# Imbalanced Brain Connectome: A Novel Link between Functional Somatic Disorders, Multiple Chemical Sensitivity, and Post-COVID patients

**DOI:** 10.1101/2025.10.22.683928

**Authors:** Shirin Haghshenas Bilehsavar, Leonardo Bonetti, Carsten Gleesborg, Sigrid Juhl Lunde, Therese Ovesen, Torben Sigsgaard, Lene Vase, Morten L. Kringelbach, Anders Rosengaard, Per Fink, Arne Møller, Lise Kirstine Gormsen

**Affiliations:** Functional Disorders, Department of Clinical Medicine, Aarhus University, Aarhus, Denmark; Center of Functionally Integrative Neuroscience, Aarhus University, Aarhus, Denmark; Department of Psychology and Behavioural Sciences, School of Business and Social Sciences, Aarhus University, Aarhus, Denmark; Department of Public Health – Research Section of Environmental and Occupational Medicine, Aarhus University, Aarhus, Denmark; Department of Otorhinolaryngology, Head and Neck Surgery, Gødstrup Hospital, Herning, Denmark; Department of Electrical and Computer Technology, Aarhus University, Aarhus, Denmark; Center for Music in the Brain, Department of Clinical Medicine, Aarhus University & The Royal Academy of Music, Aarhus/Aalborg, Denmark; Centre for Eudaimonia and Human Flourishing, Linacre College, University of Oxford, Oxford, United Kingdom; Department of Psychiatry, University of Oxford, Oxford, United Kingdom; Department of Functional Disorders, Aarhus University Hospital, Aarhus, Denmark

## Abstract

This study aimed to investigate and assess shared patterns of structural brain connectivity across three clinical populations. Despite significant advancements in neuroimaging, structural connectivity alterations have been underexplored in functional somatic disorders (FSDs), multiple chemical sensitivity (MCS), and post-COVID-19 conditions (PC). Clinical observations suggest that olfactory symptoms may represent a missing link between these three patient groups. Using diffusion-weighted MRI and voxel-based probabilistic tractography, structural connectivity was analysed in 57 females (MCS: 16, FSD: 15, PC: 11, and 15 matched healthy controls). Independent-sample t-tests (with FDR correction) and permutation testing were applied to assess group-level differences, particularly in inter- and intra-hemispheric connectivity. A marked decrease in inter-hemispheric connectivity was observed in patients (70.9%) compared with controls (29.1%; *p* < 0.001). The finding was consistent across all diagnostic groups. These alterations remained stable following reanalysis and aggregation, reliably distinguishing patients from controls. No significant differences were found in intra-hemispheric connectivity across the patient groups. Additionally, out of 12 predefined interregional connections, ten showed significant alterations in at least one patient group. The findings reveal robust and consistent reduced inter-hemispheric connectivity in patients with MCS, FSD, and PC, highlighting a potentially shared neurobiological signature among the patient groups.

## Introduction

Disorders characterized by bodily symptoms not better explained by well-known medical or psychiatric conditions including functional somatic disorders (FSDs)^1,2^, multiple chemical sensitivity (MCS)^3^, and post-COVID-19^4^ share overlapping impairing symptoms^5,6^ that challenge traditional diagnostic boundaries and may have common underlying biological or psychosocial mechanisms.

Functional somatic disorders are frequent in all medical settings and encompass different speciality-specific syndromes^7,8^ including fibromyalgia^9,10^, irritable bowel syndrome^11^, chronic fatigue syndrome/myalgic encephalomyelitis^9,10^, and MCS^12^. Empirical research identifies a multisystem FSD phenotype, also called multi-organ BDS, involving cardiopulmonary, gastrointestinal, musculoskeletal, and general symptom domains^7,8^.

Multiple chemical sensitivity, categorizing under single-organ, general fatigue symptom type, or multiorgan FSDs^12^ characterized by intolerance to odours and a range of bodily symptoms attributed to low-level exposure to environmental chemicals, typically in non-toxic doses^13^. The symptoms, often involving multiple organ systems, generally subside once the external trigger is removed and reappear only upon re-exposure to it^13^. In Denmark, a 10% prevalence of FSDs was reported in 2017^14^ and cause significant impairments^15^, poor long-term treatment outcomes^16,17^, and socioeconomic costs^18,19^. Similarly, studies highlight the high prevalence of MCS in industrialized nations, ranging from 7.5% in Japan^20^ to 19% in Sweden^21^ and 0.5%-6.5% in Denmark^22^ raising concerns about public health^23^.

The WHO defines "post-COVID-19" as the presence of at least one symptom that typically appears three months after the initial infection and persists for over two months without being attributable to other medical or psychiatric conditions^4,5,24^. Post-COVID-19 has been proposed to share a similar aetiological profile with functional somatic symptom disorders due to the symptom presentation^25^. Particularly, the issue of smell and taste dysfunction has received considerable critical attention^26^. Olfactory dysfunction has been reported from 70.2% to 85.9% in the acute phase of COVID-19^27^ as well as loss of smell (37%) in the chronic phase^28^. Fatigue is reported as the most prevalent symptom (affecting 50% of individuals with self-reported post-COVID-19)^24^. Emerging evidence indicates that prolonged symptoms following COVID-19 infection may increase vulnerability to persistent somatic symptoms^29^. Notably, several studies have highlighted overlapping between persistent COVID-19 symptoms and chronic fatigue syndrome as well as functional neurological disorder (FND)^6,30^ and somatic symptom disorder^5,6^.

Most prior research has predominantly focused on epidemiological^31^, psychological^15^, genetic^32^, and immunological dimensions^33^ of FSDs and MCS rather than neuroscientific investigations. While brain investigations in FSDs are still in their early stages, the initial findings highlight the critical role of neuroimaging techniques in elucidating the underlying pathophysiology of FSDs^34–36^. A small number of functional neuroimaging studies on MCS have consistently identified a strong and diverse array of large-scale network alterations in individuals^37,38^, suggesting that brain function entails the synchronized integration of information across a network of specialized brain regions^37^. However, to the best of our knowledge, studies indicating structural changes in MCS are non-existent.

Taken together, the key yet unresolved questions are: (1) Do individuals with FSDs, MCS, and post-COVID exhibit significant structural brain differences compared with healthy controls, which may offer insights into the underlying pathophysiology of these conditions? (2) Is there evidence to support a shared underlying pathophysiological basis across FSDs, MCS, and post-COVID? (3) Given several smell and taste dysfunction reports in post-COVID patients, can a similar dysfunction be observed in FSDs and MCS? (4) Can the brain olfactory network serve as a reference framework in structural connectome analyses and enhance the interpretation of potential structural alterations?

This study hypothesizes that FSDs, including MCS and possibly post-COVID-19, share common complex underlying mechanisms involving the olfactory system. In addition, whole brain connectivity analyses with a particular focus on olfactory networks may represent a missing link between these three patient groups: FSD, MCS, and post-COVID.

## Materials and methods

### Participants

This study included in total 42 female patients aged 18–70 years who were well-matched in terms of educational level: 16 patients with MCS, 15 patients with FSD, 11 patients with post-COVID, and 15 healthy controls (Table 1). FSD patients were recruited from the Department of Functional Disorders at Aarhus University Hospital. MCS patients were recruited from patients’ organizations and through public advertisements. Post-COVID patients were recruited from the Post-COVID Clinics at the Department of Otorhinolaryngology, Gødstrup Hospital and the Department of Clinical Medicine, Aarhus University, while healthy controls were recruited from the general population through public advertisements. Primary screening of all participants (FSD/MCS/PC/HC) was conducted over the telephone.

**Table 1.** Demographic Characteristics of Participants (N = 57)

| Characteristics | HC(n=15) | FSD(n=15) | MCS(n=16) | PC(n=11) | p-value |
| --- | --- | --- | --- | --- | --- |
| Age (years) | 49.9 ± 10.4 | 47.8 ± 11.4 | 50.8 ± 6.6 | 50.5 ± 9.1 | ,809<sup>a</sup> |
| BMI (kg/m<sup>2</sup>) | 25.1 ± 1.8 | 25.6 ± 3.8 | 24.8 ± 5.2 | 26.1 ± 4.0 | ,838<sup>a</sup> |
| *Smoking status* |  |  |  |  |  |
| Non-smoker | 100.0% | 75.0% | 86.7% | 100.0% | ,098<sup>b</sup> |
| Smoker | 0.0% | 25.0% | 13.3% | 0.0% |  |
| *Dominant hand* |  |  |  |  |  |
| Right | 92.9% | 93.8% | 80.0% | 72.7% | ,246<sup>b</sup> |
| Left | 7.1% | - | 20.0% | 27.3% |  |
| Ambidextrous | - | 6.3% | 0.0% | - |  |
| *Education level* |  |  |  |  |  |
| Primary | - | 6.3% | 6.7% | - | ,660<sup>b</sup> |
| High school | 7.1% | 18.8% | 6.7% | - |  |
| Bachelor | 35.7% | 43.8% | 40.0% | 66.7% |  |
| Master+ | 57.1% | 31.3% | 46.7% | 33.3% |  |
| *Disease duration* |  |  |  |  |  |
| Less than 5 years | NA | 37.5% | 14.3% | 100.0% | <,001 |
| 5 to 10 years | NA | 31.3% | 14.3% | - |  |
| More than 10 years | NA | 31.3% | 71.4% | - |  |

The data are shown as the mean ± SD. HC, healthy controls; FSD, patients with functional somatic disease; MCS, patients with multiple chemical sensitivity; PC, patients with post-COVID situation; BMI, Body Mass Index; NA, not applicable.
<sup>a</sup>One-way ANOVA was used to test the difference across the four groups.
<sup>b</sup>Chi-square test was used to test the difference across the four groups.

All participants received verbal and written information about the study, and written informed consent was obtained from each participant before the experiment began. In addition, all methods were carried out in accordance with relevant guidelines and regulations and in compliance with the Declaration of Helsinki.

At the first visit, all patients had a medical and psychiatric assessment by diagnostic interviews using the Schedules for Clinical Assessment in Neuropsychiatry (SCAN) and, if necessary, a clinical examination (medical and neurological) to rule out the presence of other underlying diseases or disorders was carried out by an experienced physician who was also well trained in interview techniques. Based on the screening and diagnostic interview, patients who provided informed written and oral consent, fulfilled the inclusion criteria, and did not meet any exclusion criteria were enrolled in the study.

#### FSD group

Patients with FSD were recruited based on the multi-organ diagnostic criteria for FSD/BDS according to the SCAN interview. Eligibility required the presence of multiple physical symptoms persisting for at least 6 months, involving three or more symptoms across a minimum of three different organ systems, absence of a better medical or psychiatric explanation (i.e., differential diagnoses), and the presence of significant functional impairment.

#### MCS group

Patients with MCS were diagnosed according to the 1999 consensus and the Lacour et al. 2005 criteria using the MCS checklist^39,40^ that included: A) at least two types of trigger exposure (e.g., perfume and tar); B) at least one symptom of the CNS (e.g., headache, dizziness); C) at least one symptom of either the respiratory system (e.g., nose, mouth, and eyes were included) or skin, heart/chest, muscles/joints, bladder, and stomach. Furthermore, their condition had to have resulted in impairment of daily life, social life, and work life. The duration of the diagnosis had to be at least 6 months and should meet the general inclusion criteria of the study.

#### Post-COVID

**group:** Patients referred to the Post-COVID clinics, fulfilling the WHO criteria for post-COVID-19/long COVID-19 for at least three months were also screened based on the exclusion criteria listed below. Healthy controls had no medical or psychiatric diseases and received no medication.

General exclusion criteria for all four groups: a) Before the beginning of the study: 1) Current sinonasal illness or upper respiratory tract allergy; 2) Serious or unstable medical illness (e.g., apoplexy, Parkinson’s, Alzheimer’s disease, ischemic extremity pain, renal failure, liver failure, epilepsy, and Raynaud’s phenomenon) as confirmed by medical history and, if possible, compared to medical records; 3) Current and previous diagnosis of mania, bipolar disorder, psychosis, severe agitation, imminent deliria, current suicide risk, and alcohol or drug dependence (ICD-10) confirmed by psychiatric history and, if possible, compared to psychiatric or medical records; 4) Pregnancy and lactation; 5) MR scanner incompatibility, assessed by an MRI control questionnaire. b) After the beginning of the study: 1) MRI diagnosis of incidental pathologic findings, e.g., lesions and haemorrhages; 2) The onset of acute depression or severe anxiety; 3) Suicide risk; 4) Pregnancy and lactation; 5) The patient’s desire to withdraw from the study; 6) The patient’s inability to cooperate during the examination.

In addition, treatment with antidepressants (TCAs, SSRIs, SNRIs, MAOIs, or others), gabapentin, pregabalin, and carbamazepine had to have been stopped at least 2 weeks before visit 1. All participants underwent a telephone screening, clinical examination, two test sessions, consisting of two different assessments: Clinical and neuropsychiatric assessments and smell and taste sensitivity testing, as well as one brain MRI session.

### Clinical and neuropsychiatric assessments

All patients were evaluated by a psychiatrist and underwent both medical and psychiatric assessments, including diagnostic interviews conducted with the Schedules for Clinical Assessment in Neuropsychiatry (SCAN). In addition, all participants were assessed using the Major Depression Inventory (MDI) as shown in Table 2.

**Table 2.** Neuropsychiatric and Chemosensory Assessments of participants.

| **Neuropsychiatric assessments** | **HC(n=15)** | **FSD(n=15)** | **MCS(n=16)** | **PC(n=11)** | **p-value** |
| --- | --- | --- | --- | --- | --- |
| Smell-threshold | 6.9 ± 1.4 | 6.3 ± 1.8 | 6.9 ± 2.2 | 4.0 ± 3.2<br>* | ,005<sup>a</sup> |
| Smell-Identification | 14.5 ± 2.1 | 15.3 ± .7 | 15 ± .8 | 9.45 ± 4.0<br>* | <,001<sup>a</sup> |
| TDT score | 31.5 ± 2.4 | 30 ± 2.8 | 30 ± 3.2 | 22 ± 8.5<br>* | <,001<sup>a</sup> |
| SNOT-22 | 4.47 ± 3.54 | 37.2 ± 12.4<br>* | 25.5 ± 16.3<br>* | 30.7 ± 13.8<br>* | <,001<sup>a</sup> |
| MDI | 2.00 ± 1.3 | 18.29 ± 8.5<br>* | 11.06 ± 8.9<br>* | 9.64 ± 4.8<br>* | <,001<sup>a</sup> |

Values are presented as mean ± standard deviation, HC= Healthy control group; FSD= patients with functional somatic disease; MCS= patients with multiple chemical sensitivity; PC= patients with post-COVID situation; TDT= Taste Detection Threshold test; SNOT-22= Sino-Nasal outcome Test; MDI= Major Depression Inventory; n= number; A star within the table indicates a statistically significant difference.
<sup>a</sup>One-way ANOVA was used to test the difference across the four groups.

### Smell and taste sensitivity testing

Olfactory tests. As previously mentioned, patients with MCS and post-COVID often describe diverse olfactory disturbances. Therefore, all participants underwent assessment with the psychophysical olfactory test: Sniffin’ Sticks as well as the Taste Detection Threshold (TDT) test. The purpose of the smell and taste tests was to quantitatively assess odour and taste detection thresholds and identification abilities (Fig. 1 and Table 2).

**Fig. 1.**
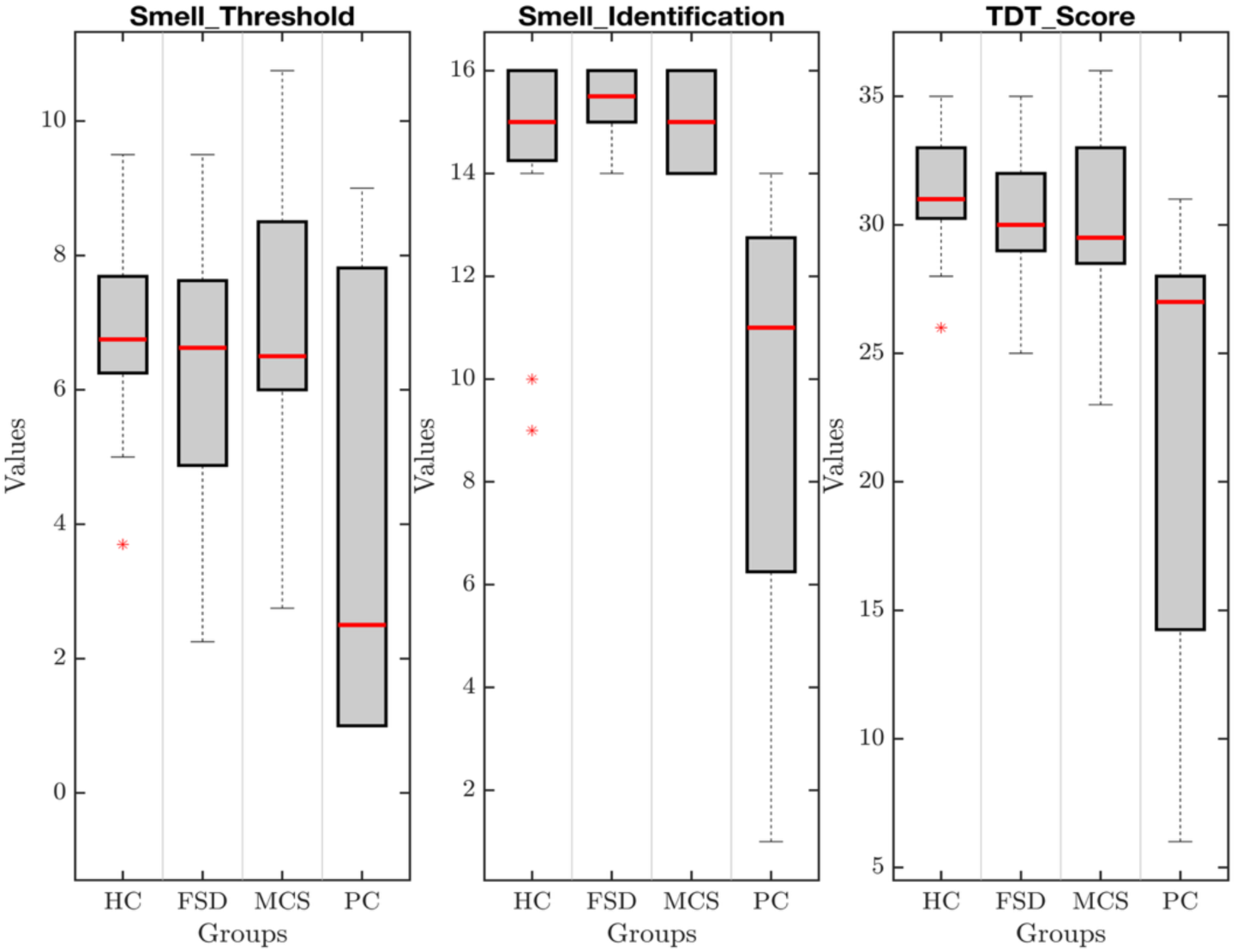
Smell and taste tests across diagnostic groups and healthy controls. Showes the distribution of Smell-Thresholding, Smell-Identification, and TDT scores across the four study groups. Each box displays the median, interquartile range, whiskers (1.5×IQR), and (*) denoting outliers. Red lines indicate the mean value for each group. The PC group shows a significant decreased scores in olfactory and taste performance compared with the other groups, while FSD and MCS groups were comparable to healthy controls. HC: Healthy Control group, FSD: Functional Somatic Disorder, MCS: Multiple Chemical Sensitivity, PC: Post-COVID.

### Brain MRI acquisition session

We employed T1 structural imaging and DTI scanning for a region-to-region reconstruction of a structural connectome in order to detect possible major neuronal pathways connecting grey matter areas. Brain scanning was accomplished through the following procedure: Whole-brain imaging was employed for all four groups, including FSD, MCS, PC, and control, using a 3T Prisma magnetic resonance imager (Siemens, Germany) with a standard head coil. For every patient, we performed 3D T1 weighted structural imaging and diffusion-weighted imaging with the following parameters; for T1 weighted sequences: Matrix size: 240×256×176; slice thickness: 1mm; repetition time (TR): 5000 ms; echo time (TE): 2.98 ms; flip angle (α): 8°, and diffusion-weighted imaging: repetition time (TR): 3300 ms; echo time (TE): 104 ms; voxel size: 2 × 2 × 2.50 mm, slice thickness of 2.50 mm, matrix size 96 × 96 ×48, and flip angle: 90°; we also used a multi-shell sequence having varied b-values.

### Preprocessing of images

In the preprocessing stage, the FDT toolbox in FSL (FMRIB Software Library, Oxford, version 5.0, www.fmrib.ox.ac.uk/fsl/) was employed, which included BET, BEDPOSTX, and PROBTRACKX. The procedure comprised head movement adjustments with FSL’s EDDY, artifact removal using the unring tool (https://bitbucket.org/reisert/unring), and FSL’s TOPUP, which estimates and makes adjustments for susceptibility artifacts. We arranged the estimation to make two fibre directions within BEDPOSTX for crossing fibres.

For brain Parcellation, the Automated Anatomical Labelling (AAL) atlas was used with 90 cortical and subcortical regions. We also used FSL’s FLIRT registration tool (FMRIB, Oxford, UK) to align data. Both DTI and the standard ICBM152 template in MNI space were registered to each subject’s native T1 space using two separate transformation matrices: one for DTI to T1 native space and another for MNI to T1 native space. These matrices were then combined to enable the transformation of the AAL atlas from MNI space into the native DTI space. (Fig. 2 illustrates the preprocessing steps and this transformation process).

**Fig. 2.**
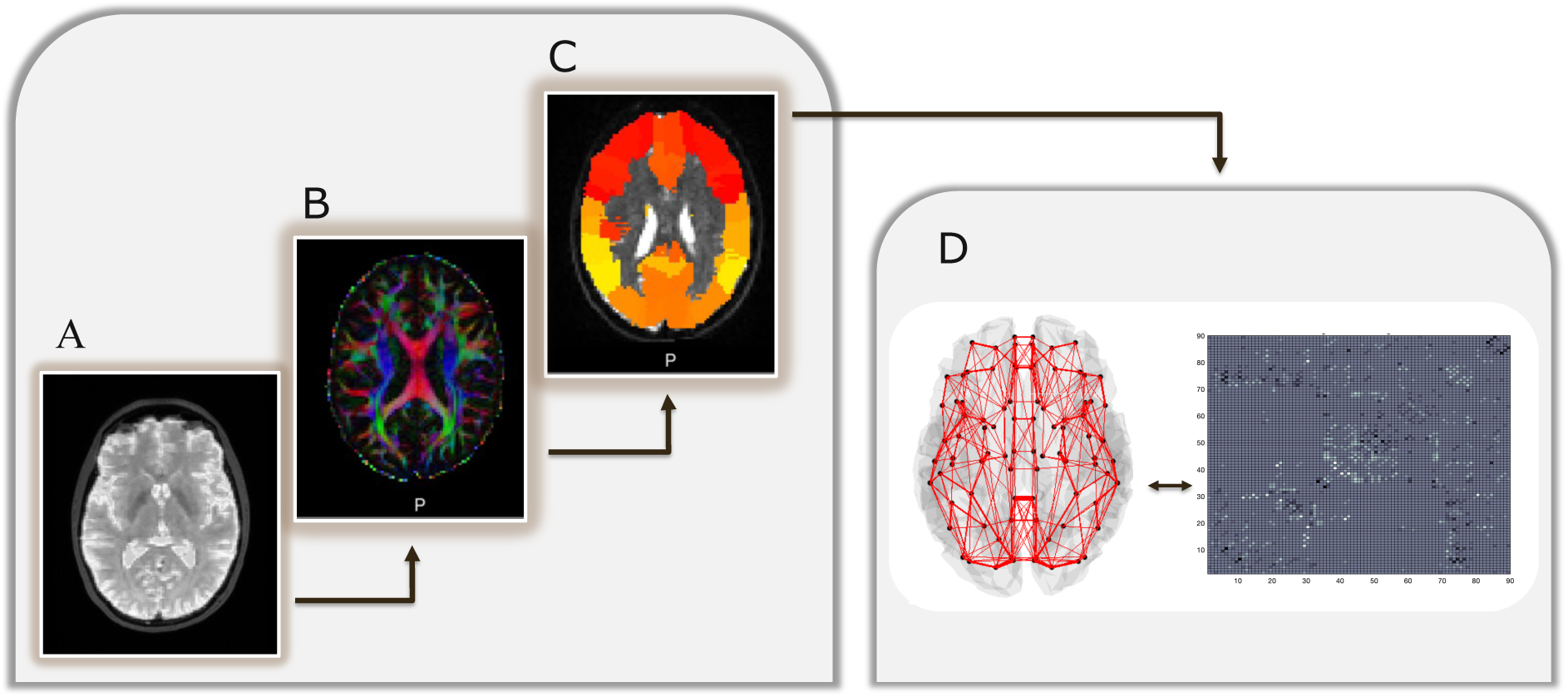
Overview for structural connectome construction. (A-B-C) Standard preprocessing pipeline showing co-registration (CR) of diffusion-weighted image (DTI-b0) to T1-weighted structural MRI (MRI-T1), transformation to standard space (MNI-T1) and computation of fractional anisotropy (FA) and probabilistic tractography in Automated Anatomical Labeling (AAL) space. (D) Final connectome generation: the whole-brain probabilistic tractography resulted in a 3D network representation (left) and corresponding connectivity matrix (right).

### Probabilistic structural connectivity

We employed voxel-level probabilistic tractography to estimate structural connectivity. Each of the 90 parcellated regions was used as a seed region, and 5000 streamlines were sampled per voxel in each region to target the remaining 89 regions. The procedure was repeated for each of the 90 regions, using them as seeds. A 90 × 90 symmetric weighted matrix was generated for each participant to model the brain’s structural network. We set a minimum threshold of 1% of the maximum connectivity strength between any two regions in a subject’s connectome to remove any spurious connections. This thresholding method was consistent across the network and scalable from voxel-level to region-level analysis.

### Group differences of structural connectivity

First, we investigated specific connections hypothesized to differ between the three patient categories and the control group. Tract selection was guided by our initial assumptions, shared clinical features (notably olfactory dysfunction), and evidence from prior neuroimaging studies. These connections were analysed using independent sample t-tests contrasting the control group versus the following patient groups: FSD, MCS, and PC. Results were considered significant only if they survived the false discovery rate (FDR) correction, with a q-value threshold of 0.0014 for FSD, 0.0189 for MCS, and 0.034 for PC patients. We selected 12 specific brain regions for focused analysis, primarily based on prior literature (Table 3). In cross-sectional neuroimaging studies, the absence of baseline data limits the ability to confidently attribute findings to the disease unlike in longitudinal cohort designs. To address this limitation and strengthen the interpretability of our results, we applied this targeted approach as a form of quality control.

**Table 3.**
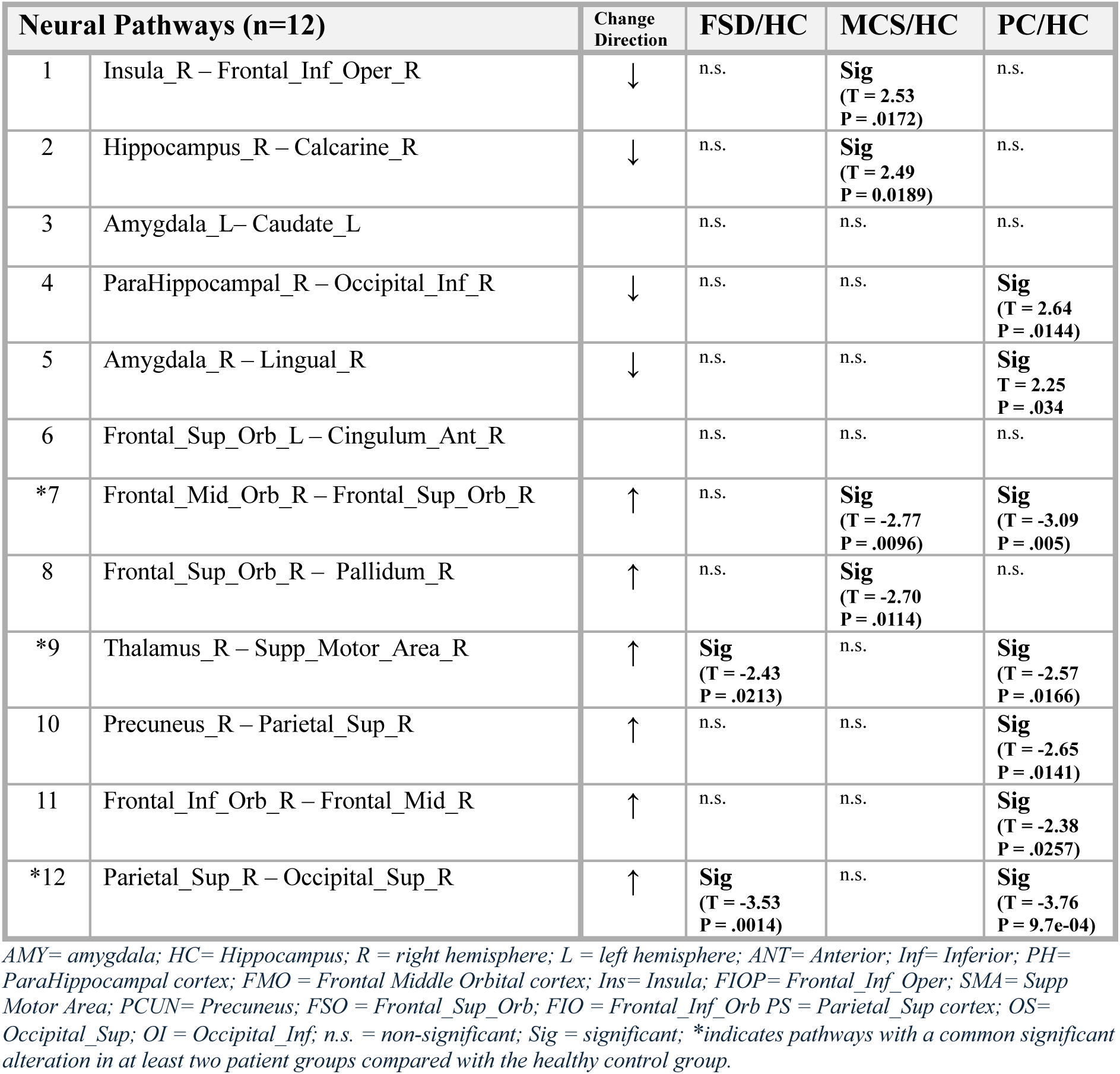
Targeted Analysis of Specific Neural Pathways.

| Neural Pathways (n=12) |  | Change Direction | FSD/HC | MCS/HC | PC/HC |
| --- | --- | --- | --- | --- | --- |
| 1 | Insula_R – Frontal_Inf_Oper_R | ↓ | n.s. | **Sig**<br>(T = 2.53<br>P = .0172) | n.s. |
| 2 | Hippocampus_R – Calcarine_R | ↓ | n.s. | **Sig**<br>(T = 2.49<br>P = 0.0189) | n.s. |
| 3 | Amygdala_L– Caudate_L |  | n.s. | n.s. | n.s. |
| 4 | ParaHippocampal_R – Occipital_Inf_R | ↓ | n.s. | n.s. | **Sig**<br>(T = 2.64<br>P = .0144) |
| 5 | Amygdala_R – Lingual_R | ↓ | n.s. | n.s. | **Sig**<br>T = 2.25<br>P = .034 |
| 6 | Frontal_Sup_Orb_L – Cingulum_Ant_R |  | n.s. | n.s. | n.s. |
| *7 | Frontal_Mid_Orb_R – Frontal_Sup_Orb_R | ↑ | n.s. | **Sig**<br>(T = -2.77<br>P = .0096) | **Sig**<br>(T = -3.09<br>P = .005) |
| 8 | Frontal_Sup_Orb_R – Pallidum_R | ↑ | n.s. | **Sig**<br>(T = -2.70<br>P = .0114) | n.s. |
| *9 | Thalamus_R – Supp_Motor_Area_R | ↑ | **Sig**<br>(T = -2.43<br>P = .0213) | n.s. | **Sig**<br>(T = -2.57<br>P = .0166) |
| 10 | Precuneus_R – Parietal_Sup_R | ↑ | n.s. | n.s. | **Sig**<br>(T = -2.65<br>P = .0141) |
| 11 | Frontal_Inf_Orb_R – Frontal_Mid_R | ↑ | n.s. | n.s. | **Sig**<br>(T = -2.38<br>P = .0257) |
| *12 | Parietal_Sup_R – Occipital_Sup_R | ↑ | **Sig**<br>(T = -3.53<br>P = .0014) | n.s. | **Sig**<br>(T = -3.76<br>P = 9.7e-04) |

AMY= amygdala; HC= Hippocampus; R = right hemisphere; L = left hemisphere; ANT= Anterior; Inf= Inferior; PH= ParaHippocampal cortex; FMO = Frontal Middle Orbital cortex; Ins= Insula; FIOP= Frontal\_Inf\_Oper; SMA= Supp Motor Area; PCUN= Precuneus; FSO = Frontal\_Sup\_Orb; FIO = Frontal\_Inf\_Orb PS = Parietal\_Sup cortex; OS= Occipital\_Sup; OI = Occipital\_Inf; n.s. = non-significant; Sig = significant; \*indicates pathways with a common significant alteration in at least two patient groups compared with the healthy control group.

**Table 4.** Supplementary Neuropsychiatric and Chemosensory Assessments Table.

| Neuropsychiatric assessments | HC(n=15) | FSD(n=15) | MCS(n=16) | PC(n=11) | p-value |
| --- | --- | --- | --- | --- | --- |
| Smell-threshold | 6.9 ± 1.4<br>6.95/6.75<br>(3.75-9.5) | 6.36 ± 1.8<br>6.36/6.5<br>(2.25-9.5)<br>.34<sup>c</sup><br>vs MCS: .74<sup>a*</sup><br>vs PC: .04<sup>a*</sup> | 6.9 ± 2.2<br>7/6.5<br>(2.75-10.75)<br>.95<sup>c</sup><br>vs FSD: .74<sup>a*</sup><br>vs PC: .009<sup>a*</sup> | 4.05 ± 3.2<br>4.05/2.5<br>(1-9)<br>.01<sup>c</sup><br>vs MCS: .009<sup>a*</sup> vs<br>FSD: .04<sup>a*</sup> | .005<sup>a</sup><br>.010<sup>a*</sup> |
| Smell-Identification | 14.5 ± 2.1<br>14.5/15<br>(9-16) | 15.3 ± .7<br>15.4/15.5<br>(14-16)<br>.15<sup>c</sup><br>vs MCS: .88<sup>a*</sup><br>vs PC: <.001<sup>a*</sup> | 15 ± .8<br>15/15<br>(14-16)<br>.42<sup>c</sup><br>vs FSD: .88<sup>a*</sup><br>vs PC: <.001<sup>a*</sup> | 9.45 ± 4.0<br>9.5/11<br>(1-14)<br>.002<sup>c</sup><br>vs MCS: <.001<sup>a*</sup><br>vs FSD: <.001<sup>a*</sup> | <.001<sup>a</sup><br><.001<sup>a*</sup> |
| TDT score | 31.5 ± 2.4<br>31.5/31.5<br>(26-35) | 30 ± 2.8<br>30.3-30<br>(25-34)<br>vs MCS: .95<sup>a*</sup><br>vs PC: <.001<sup>a*</sup> | 30 ± 3.2<br>30.1/29<br>(23-36)<br>vs FSD: .95<sup>a*</sup><br>vs PC: <.001<sup>a*</sup> | 22 ± 8.5<br>22.1/25<br>(6-31)<br>vs MCS: <.001<sup>a*</sup><br>vs FSD: <.001<sup>a*</sup> | <.001<sup>a</sup><br><.001<sup>a*</sup> |
| SNOT-22 | 4.47 ± 3.54<br>4.5/4<br>(0-13) | 37.2 ± 12.4<br>37.5/37<br>(10-59) | 25.5 ± 16.3<br>25.6/25<br>(3-62) | 30.7 ± 13.8<br>30.7/33<br>(6-46) | <.001<sup>a</sup><br>.707<sup>a*</sup> |
| MDI | 2.00 ± 1.3<br>2/2<br>(0-5) | 18.29 ± 8.5<br>18.6/18<br>(6-30)<br>.002<sup>c</sup><br>vs MCS: .03<sup>a*</sup><br>vs PC: .02<sup>a*</sup> | 11.06 ± 8.9<br>11.1/10<br>(0-26)<br><.001<sup>c</sup><br>vs FSD: .03<sup>a*</sup><br>vs PC: .89<sup>a*</sup> | 9.64 ± 4.8<br>9.6/10<br>(1-18)<br>.002<sup>c</sup><br>vs MCS: .89<sup>a*</sup><br>vs FSD: .020<sup>a*</sup> | <.001<sup>a</sup><br>.010<sup>a*</sup> |

Values are presented as mean ± standard deviation, HC: Healthy control group; FSD: patients with functional somatic disease; MCS: patients with multiple chemical sensitivity; PC: patients with post-COVID situation; TDT: Taste Detection Threshold test; SNOT-22: Sino-Nasal outcome Test; MDI: Major Depression Inventory; n: number; -/+: mean/median; (): min.-max.
<sup>a</sup>One-way ANOVA was used to test the difference across the four groups.
<sup>a\*</sup>One-way ANOVA was used to test the difference across the three patient groups.
<sup>c</sup>Two-sample t-test was used to compare the differences in tests scores between the HC and FSD, MCS, PC.

Second, we applied a permutation test (Maris and Oostenveld)^41^ to determine whether patients and controls exhibited different levels of inter-hemispheric and intra-hemispheric connectivity. This analysis was conducted for all patients together, as there was no hypothesis suggesting that they would differ across specific diseases; instead, such differences represent a common trait among these diverse patients. Initially, independent sample t-tests were performed on the original data to compare controls and patients for each connection. The underlying assumption was that if no true difference existed, the proportion of weaker and stronger connections in patients relative to controls would fluctuate randomly around 50%. However, if a systematic deviation from this distribution was observed, it would indicate a difference between the two groups. To test this, we performed 1,000 permutations of participant labels (patients and controls), maintaining the original group sizes. For each permutation, we recalculated the statistical values for each connection. We determined the percentage of weaker connections in patients compared with controls for both inter-hemispheric and intra-hemispheric connections. This process generated two reference distributions that were compared to the results from the original data. A difference in the original data was considered statistically significant if it exceeded the results from the permuted data in at least 99.9% of cases (permutation p-value < .001) (Fig. 4).

## Results

### Demographics

There were no differences in demographics between the groups including socioeconomic and educational levels. As expected, the post-COVID group had significantly shorter disease duration compared with the typically long-standing course of functional somatic disorders (Table 1).

### Clinical and neuropsychiatric assessments

Major depression was excluded in all participants across the four groups based on SCAN interviews using ICD-10 criteria; nevertheless, FSD patients showed higher MDI scores than both MCS and post-COVID groups, possibly reflecting subclinical depressive features. As expected, the Sino-Nasal Outcome Test (SNOT) scores were higher but not significantly in the patient groups than in controls (Table 2). Details are provided in supplementary Table 4.

### Smell and taste sensitivity testing

Both the threshold (*p* = .005) and identification (*p* < .001) measures demonstrated significant group differences. Compared with controls, the post-COVID group had significantly lower scores on both measures, whereas FSD and MCS did not differ from controls. Moreover, both patient groups scored higher than the post-COVID group. (Table 2) (Supplementary Table 4). Notably, the differences in the Identification subtest of PC group were more pronounced than in the threshold subtest. Although both domains were impaired in the PC group, the lower p-values observed in the Identification test suggest that PC patients experience greater difficulty in odour identification than in basic odour detection (Fig. 1).

A similar trend was observed in the Taste Detection Threshold (TDT) test, where the PC group differed significantly from both the HC group and the FSD and MCS groups, further supporting the presence of sensory impairment specific to the PC group. Results are summarized in Table 2 and illustrated in Fig. 1.

Due to intense discomfort, most participants in the MCS group and a few in the post-COVID group were unable to complete the Discrimination subtest, whereas all participants successfully completed the threshold and identification subtests.

### Brain images data analyses

We observed a robust intragroup consistency in the estimation of whole-brain structural connectivity across all three patient groups as well as the healthy control group, indicating minimal risk of artifacts such as head movement. These findings suggest a high level of reliability in our results.

### Targeted analysis of specific neural pathways

To examine specific connections hypothesized to differ, as described in the Methods section, we selected 12 neural pathways for detailed analysis in each patient group relative to controls (Table 3). Following independent-sample t-tests (with FDR correction), 10 of these pathways showed significant differences distributed across various patterns in each group (Table 3; Fig. 3). Among these 10 connections, three connections were shared by the three patient groups. In particular, connectivity between the right medial orbitofrontal cortex as well as the right superior orbitofrontal cortex was increased in both the MCS and post-COVID groups. Additionally, connectivity between the right thalamus and right superior motor area, as well as between the right superior parietal cortex and right superior occipital cortex, was increased in each of the post-COVID and FSD groups compared to controls (Table 3).

**Fig. 3.**
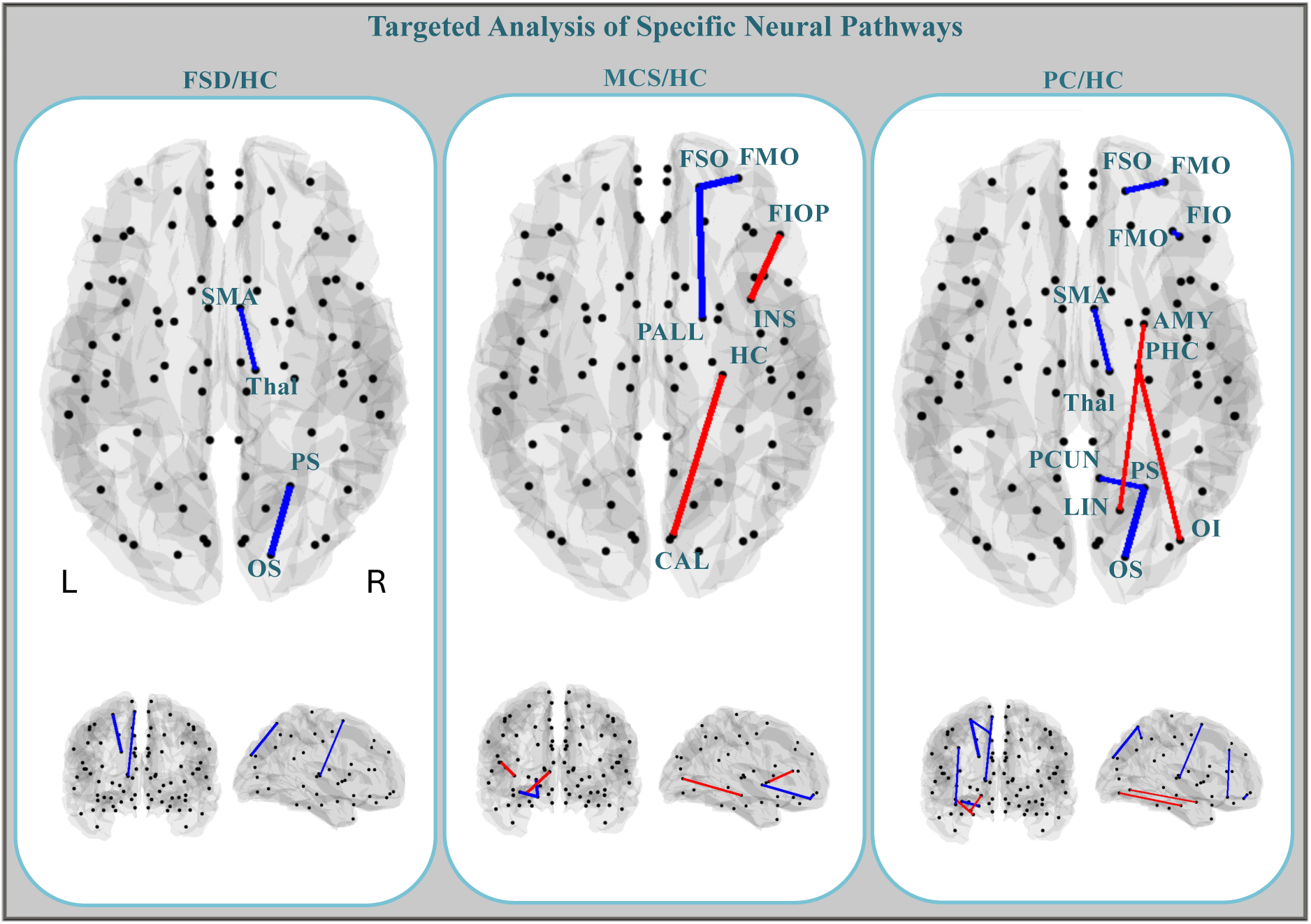
Targeted Analysis of Specific Neural Pathways. Details of the alteration in the 12 selected connectivity tracts in each patient group as compared to control group. Increased connectivity in the patient group is colour coded in blue, while decreases are coloured in red. AMY= amygdala; HC= Hippocampus; R = right hemisphere; L = left hemisphere; ANT= Anterior; Inf= Inferior; PH= ParaHippocampal cortex; FMO = Frontal Middle Orbital cortex; Ins= Insula; FIOP= Frontal_Inf_Oper; SMA= Supp Motor Area; PCUN= Precuneus; FSO = Frontal_Sup_Orb; FIO = Frontal_Inf_Orb PS = Parietal_Sup cortex; OS= Occipital_Sup; OI = Occipital_Inf.

### Comprehensive inter- and intra-hemispheric connectivity testing

In this section, we aimed to assess whether patients and controls exhibited differential levels of inter-hemispheric and intra-hemispheric connectivity using a permutation test (Maris & Oostenveld). A significant difference was observed in inter-hemispheric connectivity between controls and patients. Specifically, patients exhibited decreased connections in 70.91% of cases, whereas controls demonstrated decreased connections only in 29.08% of cases (p < .001) (Fig. 4). This finding was consistently observed across all groups, indicating a generalized decrease in interhemispheric structural integration among individuals with FSD, MCS, and PC. However, the post-COVID group showed a less pronounced reduction in interhemispheric connectivity (63.80%) compared with the FSD (71.50%) and MCS (71.90%) groups (Fig. 4).

**Fig. 4.**
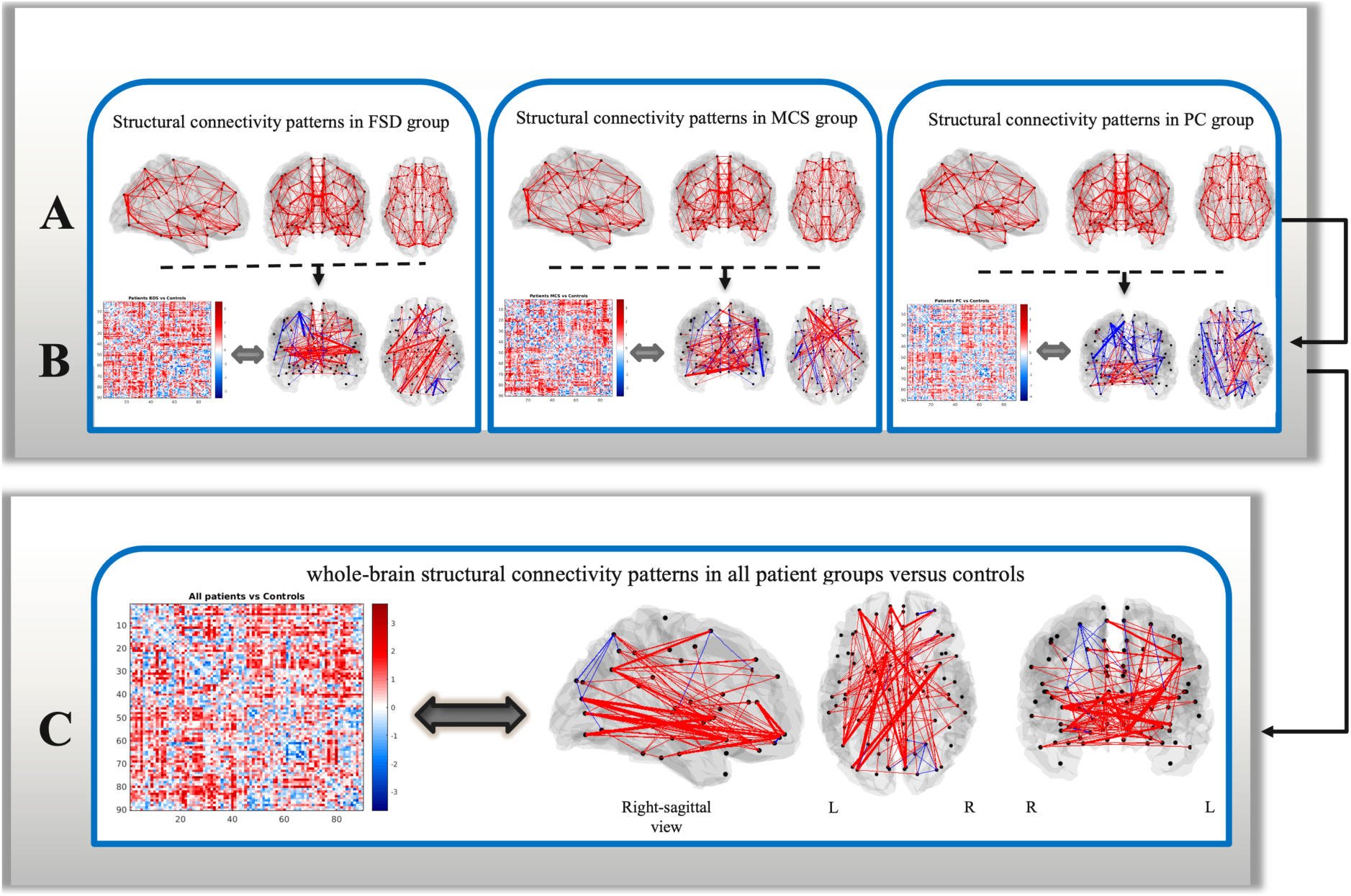
(A) Connectivity maps derived from probabilistic tractography for each single group. (B) Differences between healthy controls (HC) and patients with FSD, MCS, and PC, in group-level connectivity maps showing connections derived from probabilistic tractography. This line also includes connectivity matrices showing group-wise differences, including FSD, MCS, and post-COVID vs healthy controls (C) Whole-brain structural connectivity differences between healthy controls and all patients combined, including connectivity matrices. Blue lines denote higher connectivity for the patient groups, whereas the red lines represent lower connectivity for the patient groups. Lines are scaled according to differences in absolute connectivity.

The analysis also revealed no significant differences in intra-hemispheric connectivity within the left hemisphere (controls: 54.34% stronger connections; patients: 45.65% stronger connections) or the right hemisphere (controls: 53.23% stronger connections; patients: 46.76% stronger connections). Consequently, intra-hemispheric connectivity did not significantly differ between patients and controls (Fig. 4).

## Discussion

In this study, we demonstrate that patients with FSD, MCS, and post-COVID exhibited significantly reduced inter-hemispheric connectivity compared with controls, suggesting the potential role of altered brain structure as a relevant biological marker of the diseases (Fig. 4). In addition, the decreased interhemispheric connectivity was less pronounced in the post-COVID group than in the FSD and MCS groups, suggesting that large-scale network disruption along long-range interhemispheric pathways may be more prominent in the chronic and established conditions of FSD and MCS compared with the more recent onset observed in PC (Fig. 4).

Moreover, we show how targeted connectivities exhibit shared alterations in FSD, MCS, and post-COVID (Table 3). These findings may reflect an underlying neurobiological remodulating that contributes to the manifestation of associated symptoms in these patient groups. Of note, while a few studies have investigated structural alterations in functional somatic disorders (FSDs) using diffusion tensor imaging (DTI)^35,42^, the present study, to our knowledge represents the first structural neuroimaging investigation employing DTI specifically in a population with multiple chemical sensitivity (MCS).

In this study, we investigated voxel-level structural patterns of white matter pathways connecting grey matter regions by applying whole-brain probabilistic tractography to diffusion MRI data under strictly controlled resting-state conditions and matched environmental factors across all three groups.

### Whole-brain connectivity

We observed that the structural brain networks in three patient groups exhibited decreased inter-hemispheric structural connectivity compared with healthy controls (Fig. 4). This decrease was not limited to a specific patient group. Still, it was consistently present across all groups, indicating a generalized alteration in inter-hemispheric structural integration in individuals with FSD, MCS, and PC. In contrast, no significant alterations were detected in intra-hemispheric structural connectivity across the three patient groups.

Brain connectivity analyses make increasingly use of the concept of the "small-world brain network "^43^. This refers to the brain’s capacity to achieve high efficiency at minimal cost. In such a network, short and efficient connections, presumably less costly than long-range connections, are favoured to support optimal processing^44^. Disruption of this balance or the formation of new but inefficient connections, as observed in the patients participating in this study, may underlie changes in cognitive function or behavioural alterations^44^. Although the precise estimation of small-world metrics falls outside the scope of this study, in the absence of other graph-derived measures such as the clustering coefficient or the small-world index, the observed decrease in long-pathway efficiency connecting distant nodes (decreased inter-hemispheric connectivity) may indicate a deviation from the optimal small-world architecture and thus potentially explain the symptoms of the patients (Fig. 4). Coherently, emerging evidence indicates that alterations in small-world network topology may underlie a range of other neurological and psychiatric disorders, including schizophrenia^45^, post-traumatic stress disorder (PTSD)^46^, major depressive disorder (MDD)^47^, Bipolar mood disorders^48^, Parkinson’s disease (PD)^49^, and Alzheimer’s disease^50^. These findings are primarily supported by neuroimaging studies that consistently reveal deviations in small-world network properties among affected individuals. Such alterations are likely associated with greater metabolic and computational demands and may reflect the development of novel but inefficient neural pathways. These connectivity changes could underlie the bodily symptoms in these three patient groups.

Although the greater reduction in interhemispheric connectivity may reflect enduring structural alterations associated with chronic conditions such as FSD and MCS, this common pattern of alteration might also develop gradually over time in individuals with post-COVID condition. An alternative interpretation is that the observed decreased long-range interhemispheric connectivity, particularly within the PC group, could reflect a pre-existing vulnerability. The stability of structural connections over time may support this possibility, suggesting that certain individuals are predisposed to developing post-COVID symptoms. Accordingly, these changes may not be attributable solely to the effects of SARS-CoV-2 infection. Nevertheless, larger datasets directly comparing patients with functional somatic disorders and those with post-COVID are required to validate this interpretation.

### Specific connectivity traits

Our findings indicate that most of the targeted connectivity tracts were altered in patient groups compared with healthy controls. The selection of the 12 pathways was guided by shared clinical manifestations, particularly olfactory dysfunction, together with insights from prior neuroimaging literature (see below). Given the limited availability of structural imaging studies, particularly in MCS, we adopted a pragmatic approach, prioritizing connections associated with common symptoms in these conditions and drawing primarily on evidence from post-COVID cohorts, where structural data are relatively more available. Among the 12 targeted pathways, 10 demonstrated significant group differences (Table 3). Notably, three of these altered connections were consistently observed across at least two patient groups, suggesting the presence of potential shared neural substrates underlying these conditions (Table 3 and Fig. 3). The post-COVID group exhibited the greatest number of altered structural connections relative to healthy controls (seven of 10), exceeding those observed in the FSD and MCS groups (Table 3). The greater number of altered targeted connections observed in PC may represent an early or dynamic phase of selective network reorganization, potentially reflecting adaptive or compensatory neuroplastic mechanisms in PC group. In parallel, odour and taste detection thresholds as well as identification abilities were significantly impaired in the post-COVID group according to the smell and taste tests (Table 2). Collectively, these findings could suggest a higher degree of sensory impairment involving short-range neural pathways in post-COVID patients, whereas the FSD and MCS groups did not show significant differences in olfactory or gustatory performance compared with controls. Moreover, the convergence between connectivity alterations and reduced olfactory function may further support the reliability and validity of these observations.

Gwenaëlle Douaud’s study suggested that SARS-CoV-2 may initiate damage in primary olfactory regions, leading to secondary degeneration of the orbitofrontal cortex (OFC)^51^. Although our findings appear to contrast with previous reports that showed impairments in specific regions, they may in fact complement them. We observed increased connectivity between the medial and superior OFC in post-COVID patients compared with healthy controls, with a similar trend in the MCS group. This dissociation between regional impairments and connectivity may indicate a compensatory mechanism, whereby enhanced connectivity provides additional support to structurally affected areas.

The orbitofrontal cortex (OFC), including its medial and superior subdivisions, is a key hub in the olfactory system^52^, functionally connected to all four primary olfactory regions^52,53^. Notably, Dahmani et al. demonstrated a strong association between odour identification and spatial memory performance, which may help explain the significant decline in odour identification observed in the post-COVID group and highlight the importance of medial OFC connectivity in olfactory recognition^54^

Beyond the inter-orbitofrontal cortex, additional connections within the olfactory network were specifically examined, including those involving the insula^36,51^ amygdala^51^ hippocampus^55^ parahippocampus^56^ as well as other frontal and limbic nodes such as the inferior frontal gyrus, caudate, calcarine cortex, and occipital regions. For instance, in the MCS group, we observed decreased connectivity between the right insula and the inferior frontal operculum, as taste-related processing regions^57^. By contrast, no comparable change in this connection was detected in the post-COVID group. Nevertheless, in agreement with the observations of Esposito et al. (2021), it remains conceivable that structural remodulations could develop in post-COVID patients over a longer timeframe^58^. The same trend has been observed in connections between the right hippocampus and calcarine, further reinforcing that the connectivity between such regions is impaired in patients.

Such structural changes could represent a brain response of the CNS to a lack of sensory stimulation. In MCS patients, where long-term avoidance of odour exposure is common, the brain may undergo reorganization of the olfactory network as an adaptive mechanism aimed at stabilizing function despite the impairment. The absence of significant differences in olfactory threshold and identification scores among MCS patients may initially appear inconsistent. However, this does not necessarily rule out the presence of olfactory dysfunction in this group. The extent to which the Sniffin’ Sticks test can comprehensively assess the complexity of olfactory system function remains uncertain^59^, and in fact, the continuation of the discrimination subtest was not feasible in this study due to the heightened sensitivity and intolerance of the MCS patients. Importantly, our imaging findings nonetheless revealed structural alterations in brain connectivity patterns among affected patients, providing convergent evidence for underlying pathogenesis despite the lack of clear behavioural differences.

While a detailed discussion of individual neural pathways is beyond the aims of the present work, the observed findings can be framed within the interplay between small-world network topology and the concept of neural reserve. Small-world architecture, which balances local specialization with global integration, is thought to provide the structural basis (Bullmore & Sporns, 2009)^60^ for neural reserve, the brain’s capacity to flexibly compensate for pathology^61^. Although these concepts are distinct, our results suggest that the reduced inter-hemispheric connectivity as well as altered targeted connectivity tracts observed in FSD, MCS, and post-COVID patients may represent not only disrupted network organization but also varying degrees of preserved or impaired compensatory mechanisms. These findings appear to reflect underlying neurobiological reorganization, which may contribute to the manifestation of associated symptoms in these patient groups. This interpretation highlights the efficacy and flexibility of both whole-brain and specific connectivity tracts as potential substrates underlying resilience or vulnerability in patients with FSD, MCS, and post-COVID conditions.

## Limitations

The relatively modest sample size of this study limits the generalizability of our findings, and larger, more diverse cohorts are needed to confirm the observed alterations. Furthermore, our analyses focused on 12 predefined pathways informed by prior literature and clinical symptoms; while pragmatic, this approach may have overlooked other relevant regions or networks. The Sniffin’ Sticks test, although widely used, may not fully capture the complexity of olfactory processing, and the discrimination subtest could not be completed by all patients due to symptom intolerance, potentially reducing sensitivity. Future longitudinal studies with larger datasets, multimodal imaging, and more comprehensive olfactory assessments will be necessary to determine whether these alterations reflect transient adaptations or stable neural traits.

## Conclusions and future perspectives

This study identified consistent alterations in structural connectivity across patients with functional somatic disorders, multiple chemical sensitivity, and post-COVID with reduced inter-hemispheric connectivity emerging as a robust and convergent finding. The overlap of alterations across a limited set of targeted interregional connections among the three patient groups further reinforces the reliability of these results. Collectively, these findings suggest inefficient and reorganized long-range pathways that may constitute a shared pathophysiological substrate underlying these conditions. When interpreted within the broader framework of small-world network topology and neural reserve, the observed alterations indicate reduced inter-hemispheric connectivity and modified targeted connectivity tracts as potential markers of both disrupted network organization and variable compensatory mechanisms. Given the scarcity of structural imaging studies in functional somatic disorders, future longitudinal investigations in larger and more diverse cohorts will be essential to determine whether these alterations represent pre-existing vulnerabilities or consequences of disease processes and to clarify their role in the manifestation and progression of symptoms.

## Ethical considerations

Ethical approval for this study was obtained from the Central Denmark Region Committees on Health Research Ethics, under reference number 1-10-72-273-20, and preregistration was done at clinical.gov with the identifier NCT04935307. Further information concerning the ethical approval and the procedure for obtaining informed consent is described in the Methods section.

## Data availability

The datasets produced and analysed in this study are not publicly available in order to protect participants’ privacy; however, they are available from the corresponding author upon reasonable request.

